# Creating reproducible pharmacogenomic analysis pipelines

**DOI:** 10.1101/614560

**Authors:** Anthony Mammoliti, Petr Smirnov, Zhaleh Safikhani, Wail Ba-Alawi, Benjamin Haibe-Kains

**Affiliations:** Princess Margaret Cancer Centre, University Health Network, Toronto, Ontario, Canada; Department of Medical Biophysics, University of Toronto, Toronto, Ontario, Canada; Department of Computer Science, University of Toronto, Toronto, Ontario, Canada; Ontario Institute of Cancer Research, Toronto, Ontario, Canada; Vector Institute for Artificial Intelligence, Toronto, Ontario, Canada

**Author notes:** Corresponding author: Benjamin Haibe-Kains, Princess Margaret Cancer Centre, University Health Network, Toronto, Ontario M5G 2C4 Canada.

## Abstract

The field of Pharmacogenomics presents great challenges for researchers that are willing to make their studies reproducible and shareable. This is attributed to the generation of large volumes of high-throughput multimodal data, and the lack of standardized workflows that are robust, scalable, and flexible to perform large-scale analyses. To address this issue, we developed pharmacogenomic workflows in the Common Workflow Language to process two breast cancer datasets in a reproducible and transparent manner. Our pipelines combine both pharmacological and molecular profiles into a portable data object that can be used for future analyses in cancer research. Our data objects and workflows are shared on Harvard Dataverse and Code Ocean where they have been assigned a unique Digital Object Identifier, providing a level of data provenance and a persistent location to access and share our data with the community.

## INTRODUCTION

With the advances of high-throughput technologies in biomedicine, the volume of data has drastically increased in the last decade across scientific disciplines ^1^. This influx of data has provided researchers with the ability to discover and utilize data of various types and structural characteristics that aid in carrying out leading-edge research. However, when heterogeneous and multimodal data types are produced in large quantities, the data become much more complex to process, making conventional computational processing methods inadequate and calling for new solutions ^2,3^. These conventional methods encompass the use of scripting languages to process this data lacking (*i*) resource management capabilities (compute and memory); (*ii*) ability to aggregate data from multiple sources; (*iii*) support for modular processing; (*iv*) ability to handle unstructured data; and (*v*) ability to transform data to be used with other tools/algorithms ^4,5^. Moreover, pipelines harnessing complicated methods for processing pharmacogenomic data may be difficult to reproduce ^6^. These methods include the use of convoluted scripts that deploy multiple genomic tools and statistical methods/algorithms to compute drug response and identify molecular features in samples ^7,8^. A challenge subsequently arises, as there becomes a plethora of pipelines for pharmacogenomic datasets that utilize different complex methods, which all aim to perform the same goal, but will yield different results ^9^. These limitations hinders scalability and prevent from realizing the full potential of pharmacogenomic data generated by drug screening facilities worldwide. There is therefore a need for the development of more sophisticated computational pipelines to address these issues ^10^.

To address the issues of scalability, reproducibility and standardization with processing and analyzing pharmacogenomic datasets, we created open-source processing pipelines using the Common Workflow Language (CWL), a popular data workflow language in the data science and bioinformatics community ^11^. We leveraged *PharmacoGx* within our pipelines, an R/Bioconductor package that provides computational approaches to simplify the processing and analysis of such large datasets ^12^. We pushed our CWL pipelines on the Code Ocean platform^13^, which process two large breast cancer pharmacogenomic datasets ^14–17^ and create fully documented data objects shared through a persistent, unique digital object identifier (DOI) on Harvard Dataverse ^18^. Our study demonstrates how existing computational tools and platforms can be used to standardize the processing of pharmacogenomic data in a transparent and reproducible way, and how these processing pipelines and resulting datasets can be shared with the scientific community.

### Pharmacogenomic datasets

The first dataset is the Oregon Health and Science University (OHSU) breast cancer screen generated within Dr. Joe Gray’s laboratory (GRAY) ^14,17^.The two most recent versions of the GRAY dataset were published in 2013 and 2017, where the latest update collectively includes 91 cell lines and 120 drugs, with 10,897 drug sensitivity experiments for 72 cell lines screened against 120 drugs ^14,17^. The dataset includes processed SNP (*n*=77), exon array (*n*=56), U133A expression (*n*=51), RNA-seq (*n*=54), RPPA (*n*=49), and methylation (*n*=55) profiles with the use of various technologies and processing methods (Table 1)^17^.

**Table 1:** Summary of cell line and drug curations, sensitivity experiments, and molecular profile processing for GRAY and UHNBreast datasets

|  | GRAY 2013 | GRAY 2017 | UHNBreast 2017 | UHNBreast 2019 |
| --- | --- | --- | --- | --- |
| Cell lines | 91 | 91 | 83 | 85 |
| Drugs | 89 | 120 | 4 | 8 |
| Experiments | 9413 | 10897 | 52 | 689 |
| Molecular data and processing | RNA-seq (ALEXA-Seq, TopHat, HTSeq) |  | RNA-seq (STAR, Cufflinks) |  |
|  | CNV (aroma.affymetrix, CNTools, DNACopy) |  | CNA (Illumina GenomeStudio, CNTools, DNACopy) |  |
|  | Methylation (Illumina GenomeStudio) |  | miRNA (sva, ComBat) |  |
|  | RPPA (normalization methods from MD Anderson) |  | RPPA (normalization methods from MD Anderson) |  |

The second dataset is the University Health Network (UHN) breast cancer screen (UHNBreast) with molecular and pharmacological profiles released in 2016 ^16^ and 2017 ^15^, respectively. The dataset includes processed SNP (*n* =79), RNA-seq (*n* =82), RPPA (*n* =79), and miRNA (*n*=82) (Table 1) ^16^. We provide the most recent update to UHNBreast with four new drugs (trastuzumab, olaparib, BYL719, and UNC0642), along with processed miRNA, CNA, and RPPA data for a total of 85 cell lines, 8 drugs, and 689 drug sensitivity experiments where 56 cell lines were screened against 8 drugs^16^.

The convergence of the 2017 update of GRAY and our 2019 update to UHNBreast yield an intersection of 72 cell lines and 5 drugs after curation through our pipelines (Figure 2).

#### Reproducible and transparent processing of data

Due to the scale and complexity of data that are produced through high-throughput platforms, the data processing and analysis pipelines should possess a robust and flexible infrastructure ^4,5^. It is therefore important for pipelines to support interoperability, such as where different tools can be allocated to different data ^19^. However, pipelines that are interoperable by consisting of multiple components/stages are difficult to reproduce ^20^, such as those used to process and analyze pharmacogenomic datasets due to their multimodality ^10^. To solve this issue, we developed our *PharmacoGx* pipelines in CWL, which allowed us to standardize the way we executed our multi stage processing and analysis of both breast cancer datasets in a reproducible and transparent manner (Figure 1) ^11^. Importantly, *PharmacoGx* implements the PharmacoSet (PSet) class, allowing us to create shareable R objects integrating all aspects of pharmacogenomic datasets, from the cell line and drug annotations to the molecular and pharmacological data. Each CWL pipeline is allocated a specific subroutine that is required for PSet creation, which includes curating cell and drug annotations, computing drug response, and incorporating processed molecular profiles for a given dataset (Table 2). To accomplish this in a semi-automatic fashion, we incorporated each pipeline into a CWL workflow, where *PharmacoGx* computes each stage of a pipeline and assembles their corresponding outputs into a PSet. This workflow not only transparently indicates the pipelines that are being executed, but also ensures that each pipeline is executed in the same manner if replicated, enforcing reproducibility. Because every dataset requires a different way of transforming and processing the data, due to variability in the way the data were initially shared and structured for each study, GRAY and UHNBreast possess their own CWL pipelines and workflow to accommodate for the differences ^14–17^. Because CWL is a standardized language, each pipeline must include input and output definitions, base commands, and requirements (e.g., resource, Docker). In addition, each CWL pipeline and workflow must be accompanied by a YAML (YAML Ain’t Markup Language) or JSON file, which consists of an object array that defines a class and path for each input in the respective pipelines. In order for our CWL pipelines to execute successfully, they must specify the following: hints (docker requirement to run *PharmacoGx*), inputs that declare a type and input binding position (Rscripts, annotation files, raw drug data, processed molecular data,.), outputs that declare a type and output binding (e.g, processed drug sensitivity R objects, PSets), and a base command (to run Rscript), in the specified CWL file. Therefore, in order for our CWL workflows to be fully documentented and reproducible, each pipeline must be defined as an input and possess a successful runtime independently. Having to explicitly specify these parameters required to run each pipeline, along with the inputs and outputs in CWL provides an added layer of transparency to the pipelines, as well as allowing users to have control over data provenance. One of the highlights of our CWL workflows is the recomputation of drug response data for both datasets, which include AAC (Area Above the drug-dose response Curve), IC_50_ (maximal drug concentration to achieve 50% cell growth inhibition), Hill-Slope (measurement of slope of a drug-dose response curve), E_inf_ (maximum theoretical inhibition), and EC_50_ (drug concentration for which 50% of maximum response is observed). For GRAY, we computed sensitivity profiles that include recomputed AAC, IC_50_, Hill slope values, and included published GI_50_ (concentration for 50% of maximal inhibition of cell proliferation) data for each corresponding cell line and drug combination ^14,17^. For UHNBreast, recomputation of AAC and IC_50_ was also performed, along with Hill slope, E_inf_, and EC_50_ ^15,16^.

**Figure 1:**
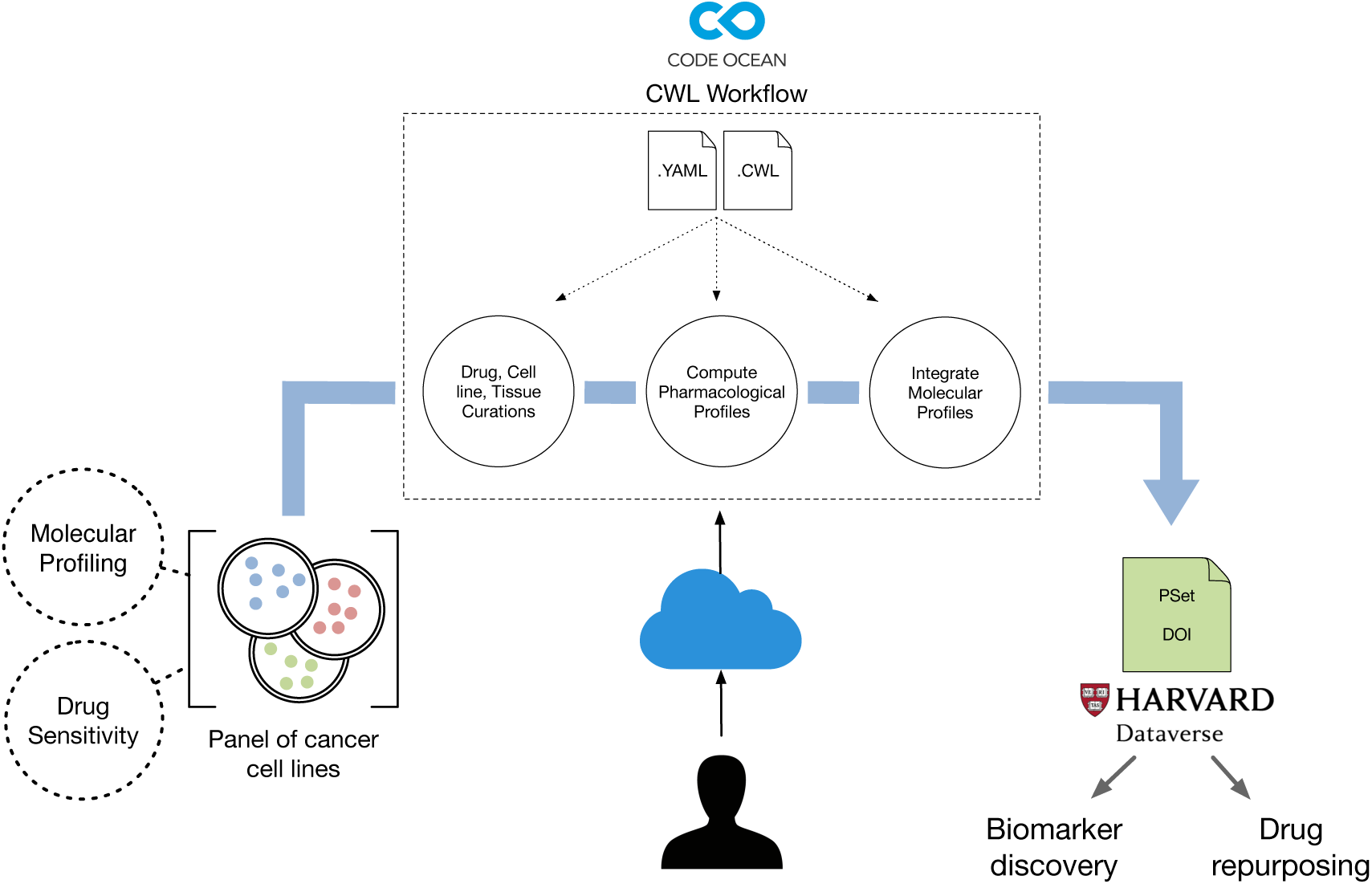
Breast cancer PharmacoSet (PSet) generation and DOI assignment through execution of a reproducible PharmacoGx CWL workflow

**Figure 2:**
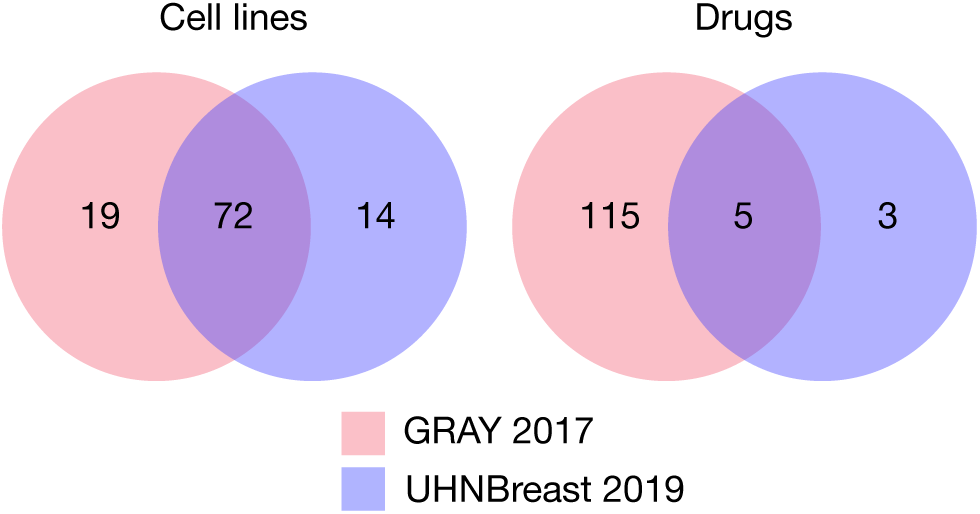
Convergence of drugs and cell lines between GRAY (2017) and UHNBreast (2019) after curation through our CWL pipelines

**Table 2:** CWL workflow pipelines and their respective data streams to produce a PharmacoSet (PSet) for GRAY and UHNBreast datasets

| CWL Pipeline | Pipeline Description | Input | Output |
| --- | --- | --- | --- |
| Cell line Curation | Curates cell lines | Cell line annotation | Curated cells line |
| Tissue Curation | Curates tissues | Cell line annotation | Curated tissues |
| Drug Curation | Curates drugs | Drug annotation | Curated drugs |
| Cell Line Info | Collects cell line metadata | Cell line metadata | Cell line metadata |
| Drug Sensitivity | Recomputes raw drug response data | Raw drug response data | Recomputed sensitivity |
| Drug Published | Collects published drug response data | Published drug response data | Published sensitivity |
| Molecular Profiles | Incorporates molecular data into ExpressionSets | Molecular profiles | ExpressionSets |
| getPSet | Creates PSet | All objects produced by each pipeline | PSet |

#### Tracking data provenance and validating pipeline integrity

Tracking data provenance with CWL can be further enhanced through the use of the provenance flag (--provenance) when executing the PSet workflows. Here, a Research Object is automatically generated, which is a directory that acts as a bundled container for all of the resources utilized and produced within our workflows, including metadata that annotates each resource ^21^. Within this object is a “data” directory that contains each input file used in the workflow with a unique and fixed checksum. We are given granular transparency across the entire workflow at every stage, as we are able to map each checksum to a respective input file and location in the “data” directory, including all of the Rscripts that were utilized within a pipeline, through a workflow metadata file that is generated. In addition to a checksum, each PSet is also assigned a Universally Unique Identifier (UUID), which provides an additional layer of provenance to accurately identify the PSet that was generated by the workflow. Moreover, this is accompanied by a provenance metadata file, which provides users with the ability to use checksums and UUID’s to accurately identify when each file was called and generated along the entire execution of a workflow. Therefore, a Research Object confirms the reproducibility of our CWL workflows and validates the PSet that was generated with a respective runtime by providing rich metadata that tracks data provenance at each stage of a workflow.

#### Harnessing Docker to create a reproducible runtime

*PharmacoGx* integrates seamlessly with CWL, as we leverage CWL’s Docker capabilities to containerize the package and run all of our pipelines in an isolated environment ^20^. Docker is a tool that allows for *PharmacoGx* to be uniformly deployed with all software dependencies, in a containerized runtime environment where all of our computations are performed and PSets are produced ^20^. The Docker container is invoked upon CWL workflow execution, where all the input files for a given pipeline become mounted into the container and all output files produced in the isolated environment are recovered into a local environment ^22,23^. Another advantage of Docker is the ability of containers to utilize and share the hardware resources of the environment it is being run in ^24^. Therefore, *PharmacoGx* deployment is not only consistent, but also portable across both cloud and high performance computing environments, as our Docker image is also publicly available through Docker Hub (https://hub.docker.com/r/bhklab/pharmacogxcwl) ^23,24^ The ability to standardize the manner in which PSets are produced through CWL, and develop an additional layer of abstraction for pipeline execution through Docker, allowed us to create and deploy reproducible and transparent pharmacogenomic pipelines that can be shared with the research community and replicated.

#### Sharing of data and pipelines

In order for a study to be computationally reproducible, data and pipelines must be well documented, uniquely identified, and easily accessible in a persistent location to other researchers ^25^. To accomplish this, we utilized the Harvard Dataverse to share our PSets for both breast cancer pharmacogenomic datasets, along with Code Ocean to share our CWL pharmacogenomic pipelines ^13,18^. Harvard Dataverse is an online data repository for transparently preserving and sharing research data with other researchers. By creating a container known as a “dataverse” within the platform, researchers are to able deposit their datasets and corresponding metadata, in an organized fashion and make them easily discoverable for others to download and share ^18^. Each dataset can be also assigned an unique DOI, which allows a dataset to possess a persistent location, as well as allow researchers to accurately identify and share a specific dataset of interest. In addition, subsequent updates (versions) to a dataset can be uploaded, with accompanying metadata that explains the update and its changes, providing a layer of data provenance to the research community.

We also transferred our reproducibility measures to the pipeline level, as we deposited and shared our CWL workflows through Code Ocean, a reproducibility platform that allows for researchers to upload, share, and run published and configured code ^13^. Data is uploaded into a “capsule”, which provides a computational environment for others to run code in the capsule, without the need to manually execute it locally with the addition of installing any dependencies ^13,26^. Moreover, code can also be assigned a persistent DOI, providing the ability to accurately share and retrieve pipelines, as well as verify the reproducibility of published results directly through the compute capsule ^13,26^. Because Code Ocean does not currently support running multi-container pipelines, and therefore our CWL workflows, we used the platform to host our workflows and raw data, provide execution instructions, and run a post-PSet analysis for biomarker discovery.

Our PSets can be found on Harvard Dataverse at the following DOI: https://doi.org/10.7910/DVN/BXIY5W. Our CWL workflows can be found on Code Ocean at the following DOI: https://doi.org/10.24433/CO.7378111.v1.

#### Utilization of PSets for Biomarker Discovery

In order to demonstrate the utilization of our PSets for cancer research, we identified ERBB2 expression as a biomarker for lapatinib in both the GRAY 2017 and UHNBreast 2019 datasets (Figure 3). To investigate this gene-drug association, we utilized processed RNA-seq expression and recomputed drug sensitivity (AAC) from each PSet. We subsequently identified cell lines from both PSets that possess gene expression and drug sensitivity for ERBB2 and lapatinib, respectively, and computed the concordance index and p-value between them. Our computed concordance index of 0.7308 and 0.6299 for GRAY and UHNBreast respectively, indicate a correlation between the effects of drug response (lapatinib) to molecular features (ERBB2 amplification) across both datasets. This analysis can be reproduced through our Code Ocean capsule.

**Figure 3:**
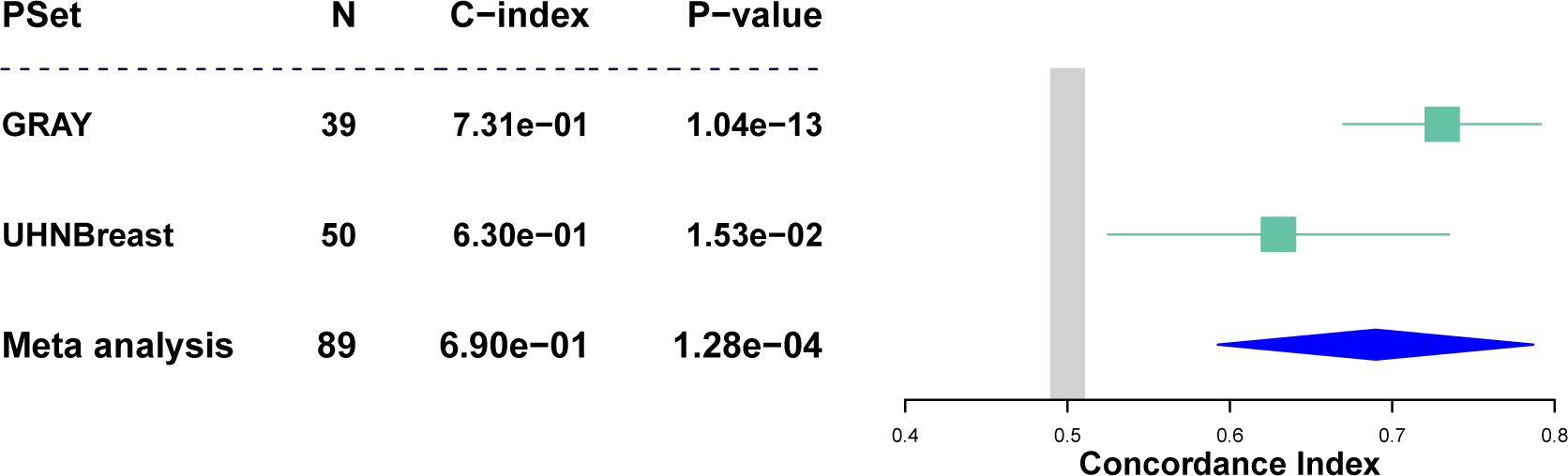
ERBB2 expression as a biomarker for lapatinib in GRAY 2017 and UHNBreast 2019. N: number of samples; C-index: concordance index calculated for respective PSet; P-value; p-value calculated for respective PSet. Meta analysis represents combined concordance index across PSets

## DISCUSSION

The utilization of CWL allows us to create and execute transparent and reproducible pharmacogenomic pipelines that can be validated and easily shared with the scientific community ^11^. The standardized architecture of the language allows users to create language-agnostic pipelines and workflows that enforce strict parameter specifications to ensure execution is consistent ^11^. In addition, users are able to incorporate Docker into their runtimes, where data ingestion, analysis, and exportation all occur in an isolated container environment that promote repeatable execution ^22,23^. Users are also able to track data provenance across the entire execution time by creating Research Objects in CWL, which validates each portion of data flow from input to output, through checksums and UUID’s ^21^. Lastly, CWL pipelines and workflows are scalable and portable across many computing environments, such as the cloud, which gives users the ability to easily share their analyses and harness a plethora of various hardware resources to successfully execute their workloads that would not be possible with using on premise resources ^23,24^. A common practice in pharmacogenomics is sharing study data as supplementary files through a journal, or online sharing platforms/repositories such as Synapse and GitHub, which was the case for both the GRAY and UHNBreast datasets ^14–17^. However, the challenge becomes assembling these data into a form that can be successfully analyzed and interpreted when shared. We were able to accomplish this in a reproducible manner by utilizing study data from a variety of sources and assembling it into a meaningful and useful form for cancer researchers, which are PSets, through CWL and *PharmacoGx.* Therefore, our pipelines form the bridge between raw pharmacogenomic data and assembly in a transparent fashion. However, our workflows do have limitations, including the inability to identify changes to pipelines, input data, and PSets, at the file level, when updates are pushed and the files are taken into an environment outside of Harvard Dataverse and Code Ocean. However, with storing our data on Harvard Dataverse and pipelines on Code Ocean with rich metadata, users will be able to retrieve any updated files on both repositories and accurately identify the exact changes to each file. In addition, CWL Research Objects provide checksums and UUID’s only after a runtime is complete, which are bound to the file name and not persistently attached to a file for use in subsequent workflow runs. Thus, if a input file is updated and re-utilized in a workflow, we must manually keep track of all checksums and UUID’s that were assigned to it by CWL over time. In the future, we hope to increase transparency and reproducibility by automating these pharmacogenomic pipelines in a manner that keeps track of all input and output data at the file level through the use of automatically generated unique identifiers that are persistent. Moreover, we hope to provide users with an interface that provides options for processing drug sensitivity and molecular profiles in a generated PSet.

## CONCLUSION

Our *PharmacoGx* CWL pipelines allow for the processing and analysis of pharmacogenomic datasets in a reproducible and transparent manner. The pipelines compute pharmacological data across many cell lines and drugs and integrate it with molecular profiles to generate a breast-cancer specific PSet object. These data objects and pipelines are made available through a persistent DOI on Harvard Dataverse and Code Ocean respectively, which provide researchers with the opportunity to transparently share and utilize them for improved biomarker discovery.

